# Follicular fluid metabolomic analysis in women with Hashimoto’s thyroiditis

**DOI:** 10.1101/2022.12.19.520992

**Authors:** Diana Caroline da Silva Bastos, Maria Izabel Chiamolera, Renata Elen Costa da Silva, Maria Do Carmo Borges De Souza, Roberto De Azevedo Antunes, Marcelo Marinho De Souza, Ana Cristina Allemand Mancebo, Patrícia Cristina Fernandes Arêas, Fernando M. Reis, Edson Guimarães Lo Turco, Flavia Fonseca Bloise, Tania Maria Ortiga-Carvalho

## Abstract

Hashimoto’s thyroiditis is an autoimmune thyroid disease characterized by hypothyroidism and a high level of anti-thyroid autoantibodies. This disease has been linked to a negative impact on female fertility, but the mechanisms are unclear. Ovarian follicular fluid appears to be the key to understanding how Hashimoto’s thyroiditis can affect fertility. Therefore, we aimed to evaluate the follicular fluid metabolic profile and its relationship with anti-thyroid autoantibody levels. For this, we collected follicular fluid from a total of 61 patients undergoing in vitro fertilization treatment, comprising 34 women with thyroid autoantibody positivity and 18 negative controls. Follicular fluid samples were analysed using metabolomics and thyroid autoantibodies were measured. Follicular fluid samples from Hashimoto’s thyroiditis patients presented 15 metabolites with higher concentrations than those in controls, which indicates five possible affected pathways: the glycerophospholipid, arachidonic acid, linoleic acid, alpha-linolenic acid, and sphingolipid metabolism pathways. These pathways are known to regulate ovarian functions. In addition, anti-thyroglobulin antibody concentrations were more than tenfold higher in women with Hashimoto’s thyroiditis than in controls, in both serum and follicular fluid. Our data showed that Hashimoto’s thyroiditis can change the metabolic profile of follicular fluid, suggesting a potential mechanistic explanation for the association of this disease with female infertility.

## 1 Introduction

Women with Hashimoto thyroiditis (HT) suffer more from miscarriage, recurrent pregnancy loss, decreased fertilization rates, and reduced numbers of good-quality embryos than euthyroid women (Monteleone *et al*., 2011; Chen *et al*., 2017; Andrisani *et al*., 2018; Medenica *et al*., 2018; Dong *et al*., 2020). HT is the most common autoimmune disease in reproductive-age women (Krassas, Poppe and Glinoer, 2010). The two autoantibodies associated with autoimmune destruction of the thyroid gland and with the clinical condition of hypothyroidism are the anti-thyroid peroxidase antibody (TPOAb) and the anti-thyroglobulin antibody (TGAb) (Hiromatsu, Satoh and Amino, 2013). Nonetheless, treatment of HT with thyroid hormone replacement does not seem to have a protective effect on fertility outcomes. HT patients with controlled thyroid hormone levels still present a low fertility rate (Rao *et al*., 2019). Thus, the impact of HT on infertility cannot be associated only with hypothyroidism.

Follicular fluid (FF) and autoantibodies seem to play a central role in the relationship between HT and female infertility. The first study to propose this idea showed the presence of these antibodies in FF, with higher levels in women with HT (Monteleone *et al*., 2011)). After this finding was reported, other studies investigated the presence of antibodies in FF and women’s fertility outcomes (Medenica *et al*., 2018; Cai *et al*., 2019). In addition to the increased levels of autoantibodies in FF, women with HT had lower oocyte fertilization rates, fewer grade A embryos, lower pregnancy rates and an increased risk of early miscarriage compared to euthyroid women (Monteleone *et al*., 2011; Medenica *et al*., 2018)).

FF originates from the union of products released by granulosa and theca cells with blood exudate (Rodgers and Irving-Rodgers, 2010)). The composition of FF is different from that found in serum (Basuino and Silveira, 2016)). FF is composed of metabolites that accumulate in the oocyte, allowing a necessary scenario for its maturation. FF contains amino acids, lipids, nucleotides, and essential factors for oocyte competence, such as hormones, cytokines, growth factors, proteins, and metabolites (Hennet and Combelles, 2012; Dumesic *et al*., 2015).

Although some research has shown high concentrations of thyroid autoantibodies in the FF of women with HT, it is still unknown whether these factors could explain how the disease affects fertility. Regarding protein expression, 49 proteins were found to be differentially expressed in FF from HT patients compared to FF from control patients (Y. Liu *et al*., 2020). This finding suggests that HT can alter the composition of FF.

To confirm the possible impact of the disease on FF composition, it is necessary to evaluate the FF metabolic profile, since the metabolic composition of FF is important for oocyte development (Hennet and Combelles, 2012). Thus, the change in the metabolic profile of FF may be the mechanism by which HT affects female fertility.

Metabolomic analysis of FF has already been performed in search of answers to changes in ovarian follicles (Lazzarino *et al*., 2021; Liang *et al*., 2021; Wang *et al*., 2021). In addition, other studies have used the technique to further investigate the pathophysiology of HT in other samples, such as blood, urine, and thyroid biopsy samples (Tsoukalas *et al*., 2019, 2020; Capitoli *et al*., 2020; J. Liu *et al*., 2020; Piras *et al*., 2021). However, applying metabolomics to understand the link between HT and changes in ovarian follicles, particularly for FF, is still an unexplored area.

The aim of this study was to evaluate the FF metabolic profile and its relationship with anti-thyroid autoantibody levels. Understanding the mechanism by which HT can affect female fertility would provide valuable insight into the complex pathophysiology of the disease and a possible starting point to develop therapies targeted towards the associated subfertility.

## 2 Materials and Methods

### 2.1 IVF patients

The study had a prospective design. All patients eligible for IVF treatment in the Fertipraxis, a reproductive medicine center located at the city of Rio de Janeiro, from 2019 to 2020 were considered potential participants. The exclusion criteria were IVF cycle cancellation or patients who had no thyroid function information. A total of 61 women undergoing IVF were divided into two groups according to the blood levels of thyroid autoantibodies (TPOAb levels > 34 IU/mL and/or TGAb levels > 115 IU/mL) or a previous diagnosis of HT. Thus, 38 euthyroid women had thyroid autoantibody positivity and 23 were negative controls. After the selection of the patients for each group, all measurements were blinded to the researchers until final analysis.

Although we selected a total of 61 patients, some parameters were evaluated in fewer patients according to the information that was available. TGAb blood tests were available for 38 patients (26 from the HT group and 12 from the control group), while TSH and free T4 were assessed in 37 participants (14 in the HT group and 23 in the control group). Regarding the fertilization rate, 34 patients in the HT group and 18 in the control group were evaluated because eight patients had all oocytes cryopreserved.

All subjects signed an informed consent form during their first medical evaluation. None of the patients were categorized by ethnicity due to the mixed genetic heterogeneity of the Brazilian population. This research was approved by the Research Ethics Committee on Research at the Maternidade Escola, Universidade Federal do Rio de Janeiro and registered on Plataforma Brasil under number 02213812.4.0000.5275.

### 2.2 FF thyroid autoantibody measurements

FF was collected during follicular aspiration and stored at −20°C for future analyses. The fluid was used to measure autoantibody levels and to perform metabolomic tests. To investigate the concentrations of TPOAb in FF, we used the Elecsys kit competition immunoassay method (ROCHE, Mannheim, Germany) following the manufacturer’s instructions. The limit of detection was 5 UI/mL. The levels of TGAb in FF were measured by the “*In house*” assay, which has been previously described and validated (Nakabashi *et al*., 2012). The limit of detection is 10 UI/mL.

### 2.3 FF metabolomic analysis

Metabolite analysis of the patients’ FF samples was performed using the liquid chromatography method coupled to an electrospray ionization mass spectrometer (LC-ESI-MS). The data obtained such as the concentration of each metabolite identified in the FF and its normalization are available in an online repository and can be found under the number: 10.17632/nm5gzmdsjh.1.

The LC–MS analyses were performed in a Nexera CL HPLC System (Shimadzu) consisting of a degasser unit, two unitary pumps, an autosampler, a column oven and a controller module. The HPLC was coupled to a triple-quadrupole LCMS-8060 CL mass spectrometer (Shimadzu). The injection volume in all cases was 5 μL, and ionization was performed *via* electrospray ionization (ESI) in positive and negative modes, depending on the analyte. The analyses were performed using a targeted approach, and the ions were assessed by multiple reaction monitoring (MRM).

The mobile phases were prepared with acetonitrile (ACN), water and formic acid (Merck) in different proportions, all of them being LC–MS grade. The results were acquired using three different methods to evaluate the following diverse classes of the molecules of interest: (I) primary metabolites (acquired from Shimadzu); (II) lipid mediators (acquired from Shimadzu); and (III) lipids and carcinogens (*in house* development). The methods are detailed in the next paragraphs.

The mobile phase for method (I) – primary metabolites – was composed of acid 0.1% (v/v) (A) and ACN + formic acid 0.1% (v/v) (B). The analyses were carried out using a PFFP (3 μm, 150 mm x 2.1 mm, Supelco) column at 40 °C. The gradient of B, at a flow of 0.25 mL/min, was as follows: time 0, 0%; 1 minute, 0%; 2 minutes, 25%; 11 minutes, 35%; 15 minutes, 95%; 20 minutes, 95%; 20.1 minutes, 0%; and 25 minutes, 0%.

Method (II)—lipid mediators—was performed using a Kinetex C8 (2.6 μm, 150 mm x 2.1 mm, Phenomenex) column at 40 °C. The mobile phase was acid 0.1% (v/v) (A) and ACN (v/v) (B), the flow was 0.40 mL/min, and the gradient for B was as follows: time 0, 10%; 5 minutes, 25%; 10 minutes, 35%; 20 minutes, 75%; 20.1 minutes, 95%; 25 minutes, 95%; 25.1 minutes, 0%; and 28 minutes, 0%.

The column and mobile phase for method (III)—lipids and carnitines—was the same as for method (II). The column oven was set at 70 °C and the initial flow was 0.45 mL/min. The gradient for B, totalling 28 minutes of analysis, was as follows: time 0, 5%; 8 minutes, 60%; 20 minutes, 80%; 21 minutes, 98%; 26 minutes, 98%; 26,1 minutes, 5%; and 28 minutes, 5%. A flow gradient was also used for this method: time 0, 0.45 mL/min; 19 minutes, 0.45 mL/min; 19.5 minutes, 0.55 mL/min; 27 minutes, 0.55 mL/min; 27.5 minutes, 0.45 mL/min; and 28 minutes, 0.45 mL/min.

The table generated after the processing of the individual metabolites was used to carry out the multivariate analysis of the data. For this, we used MetaboAnalyst 5.0 online software. Principal component analysis and partial least squares discriminant analysis (PLS-DA) were performed on the log-transformed data and standardized by Pareto scaling. Principal component analysis was used to observe grouping and sample discrepancies in general, while PLS-DA was used to maximize the variations and to guarantee the discriminatory effect of the components based on the values of the variable importance in projection (VIP).

A cross-validation test was applied to validate the method created by PLS-DA and indicated which component of the model was the one that best explained the variations in ions between the groups. Based on the PLS-DA, we selected the metabolites with the highest VIP scores from the component with the greatest discriminatory effect as potential biomarkers. We built an ROC curve for the metabolite combinations. In addition, we performed a permutation test (1000x) and calculated the prediction class probability based on our samples.

### 2.4 Pathway analysis

The study of possible affected pathways was carried out through the listing and concentration of metabolites found by PLS-DA. This information was processed by MetaboAnalyst 5.0 software in the Pathway Analysis module. The Pathway Library – Homo sapiens (hsa - KEGG organisms) was used. The Global test method was used for pathway enrichment analysis.

### 2.5 Statistical analysis

The analysis of clinical data between the groups was performed using the non parametric 2-sided Mann-Whitney U test, using SPSS (IBM SPSS Software version 22). The results obtained from the patients’ clinical data are expressed as the median (quartiles) or numbers (percentages). Furthermore, multivariate statistics were performed using MetaboAnalyst 5.0 software. Significant differences were considered when p<0.05.

## 3 RESULTS

### 3.1 Laboratory and clinical information

Table 1 summarizes the clinical and laboratory data of the HT patients and controls. The two groups had similar characteristics in the clinical evaluation, except for the duration of infertility, which was higher in the control group (p = 0.01).

**Table 1:** Clinical and laboratory information of the Control group and the Negative Hashimoto Group.

|  | Control | Hashimoto | P value |
| --- | --- | --- | --- |
| Age (years) | 36 (35-39) | 36 (32-38) | 0.423 |
| BMI (kg/m<sup>2</sup>) | 21 (20-26) | 23 (22-26) | 0.183 |
| Duration of infertility (months) | 19 (12-36) | 9 (5-14) | 0.011 |
| Previous miscarriages | 3/16 (19%) | 10/33 (30%) | 0.502 |
| Serum TSH (μIU/ml) | 2.18 (1.46-3.64) | 1.79 (2.29-2.99) | 0.882 |
| Serum FT4 (ng/dl) | 1.20 (1.05-1.32) | 1.10 (1.00-1.45) | 0.660 |

Data are expressed as the medians (quartiles) or numbers (percentages). P values refer to the 2-sided Mann–Whitney U test. BMI: body mass index; TSH: Thyroid-Stimulating Hormone; FT4: Free Thyroxine;. Control n=23. Hashimoto n=38

The two groups had similar concentrations of TSH, FT4, serum TPOAb and FF TPOAb (Table 1 and Figure 1). However, serum and FF TGAb levels were dramatically increased in the group of patients with HT (p < 0.001, Figure 1).

**Figure 1:**
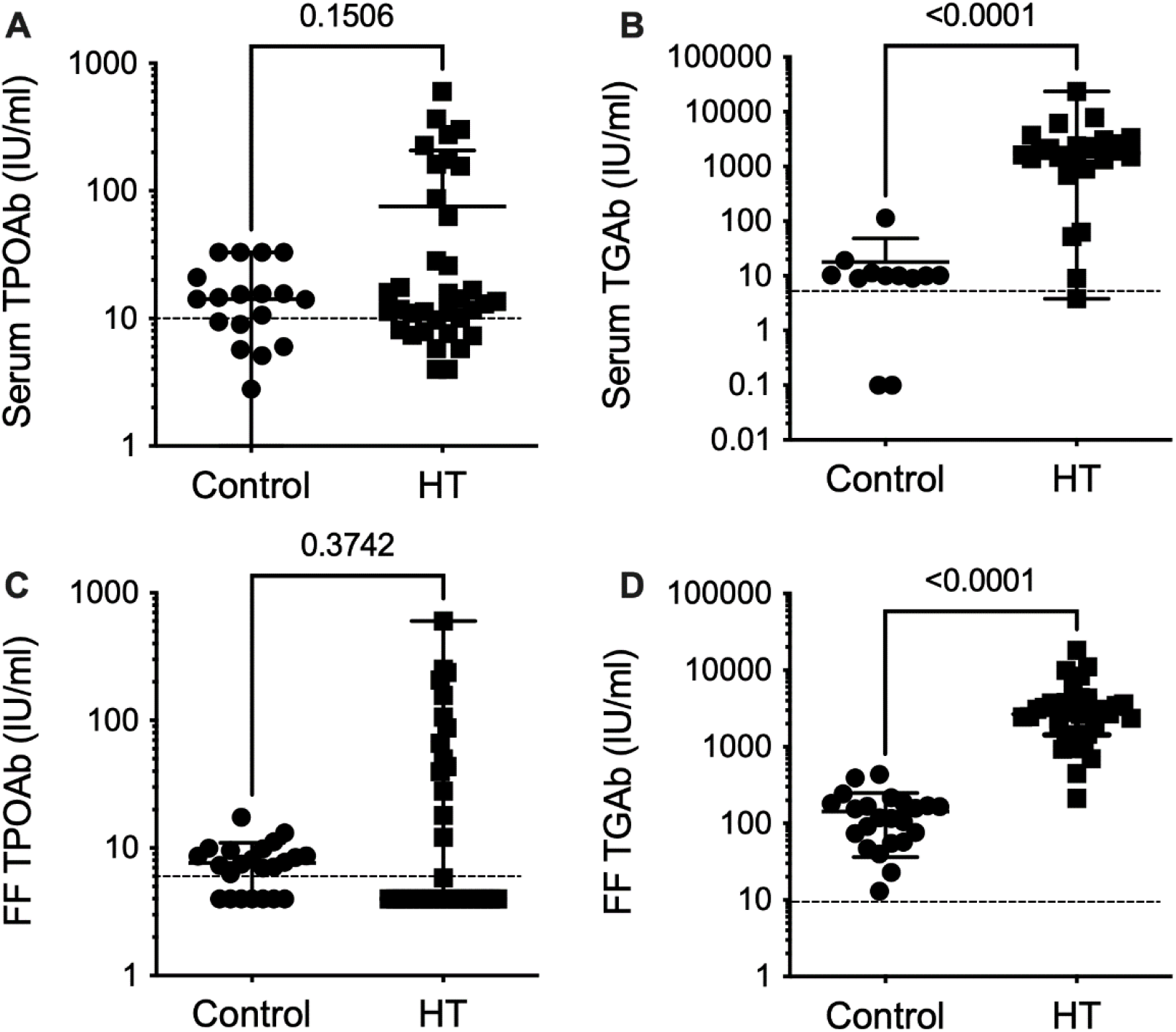
Anti-thyroid antibody levels in follicular fluid and serum samples from patients undergoing in vitro fertilization. (A) Serum anti-thyroid peroxidase (TPOAb), (B) Serum anti-thyroglobulin (TGAb), (C) Follicular fluid TPOAb, (D) Follicular fluid TGAb levels (shown for each group), Control (circle) and Hashimoto (HT square). Data are expressed as the medians (quartiles). P values refer to the 2-sided Mann–Whitney U test. Dotted line represents limit of detection of each essay. Control n=23. Hashimoto n=38.

Even though only one TGAb concentration showed a significant difference between groups (Figure 1B, D), the concentrations of TPOAb were more dispersed and skewed in the HT group (Figure 1A, C).

### 3.2 In vitro fertilization parameters

After clinical evaluation, we investigated intermediary IVF outcomes between groups (Table 2). The parameters used in the study were the number of oocytes, proportion of metaphase II oocytes, fertilization rate and number of embryos. In the analysed outcomes, we found no significant difference between the control and HT groups.

**Table 2:**
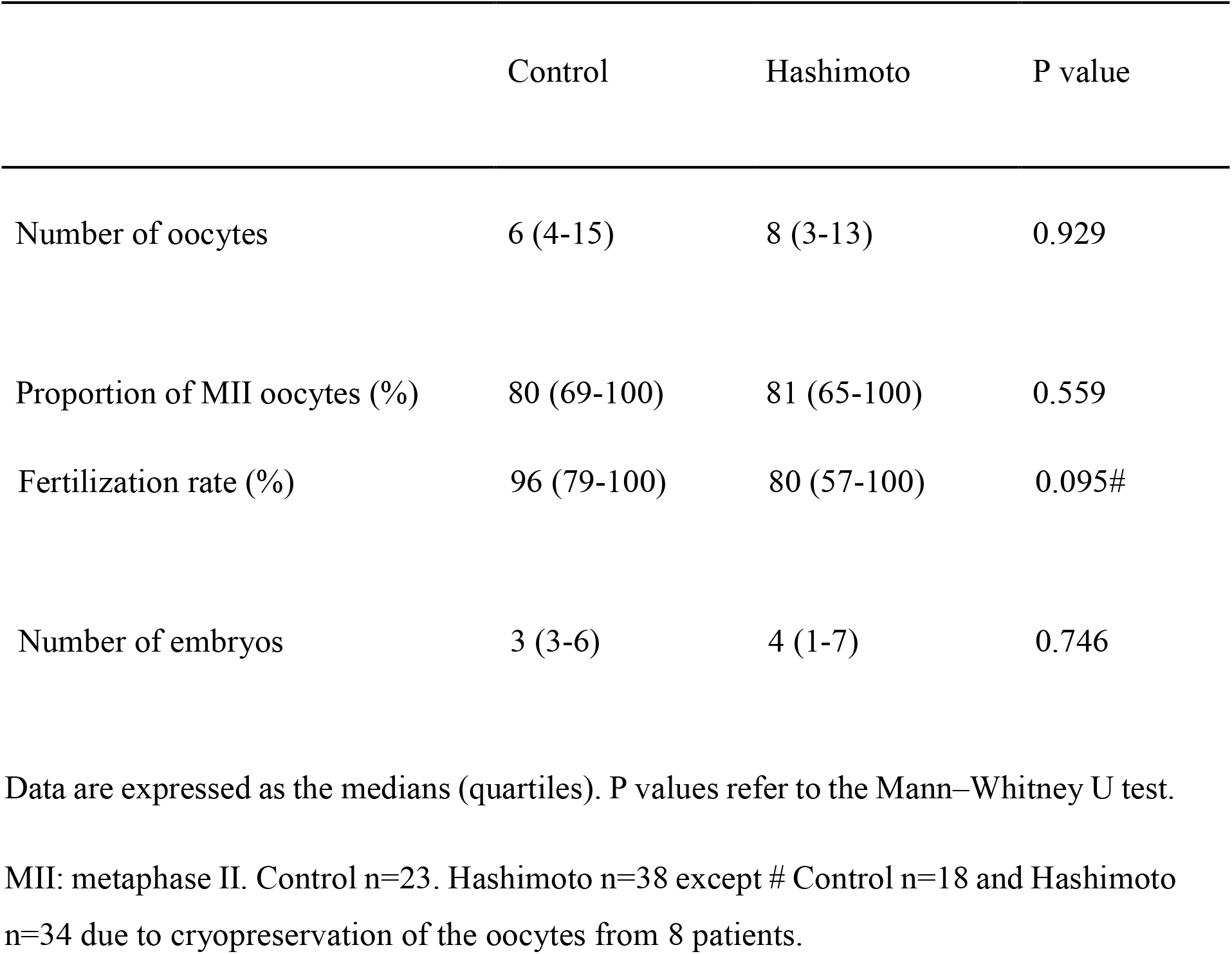
In vitro fertilization outcomes in the two groups.

### 3.3 Metabolomics of follicular fluid

In the FF metabolomic analysis, we detected and measured the concentrations of 83 metabolites. In the PLS-DA, it was possible to observe the different metabolic profiles in the FF of the two groups (Figure 2). The analysis of metabolites presents in FF showed differences in 15 metabolites between the HT and euthyroid patients (Figure 3). All 15 metabolites were found to have higher concentrations in the FF of HT patients. Among the metabolites found to be altered, there were ten phosphatidylcholines, two acylcarnitines, two sphingolipids, and one lysophosphatidylcholine. Quantitative analysis confirmed our data (Figure 4).

**Figure 2:**
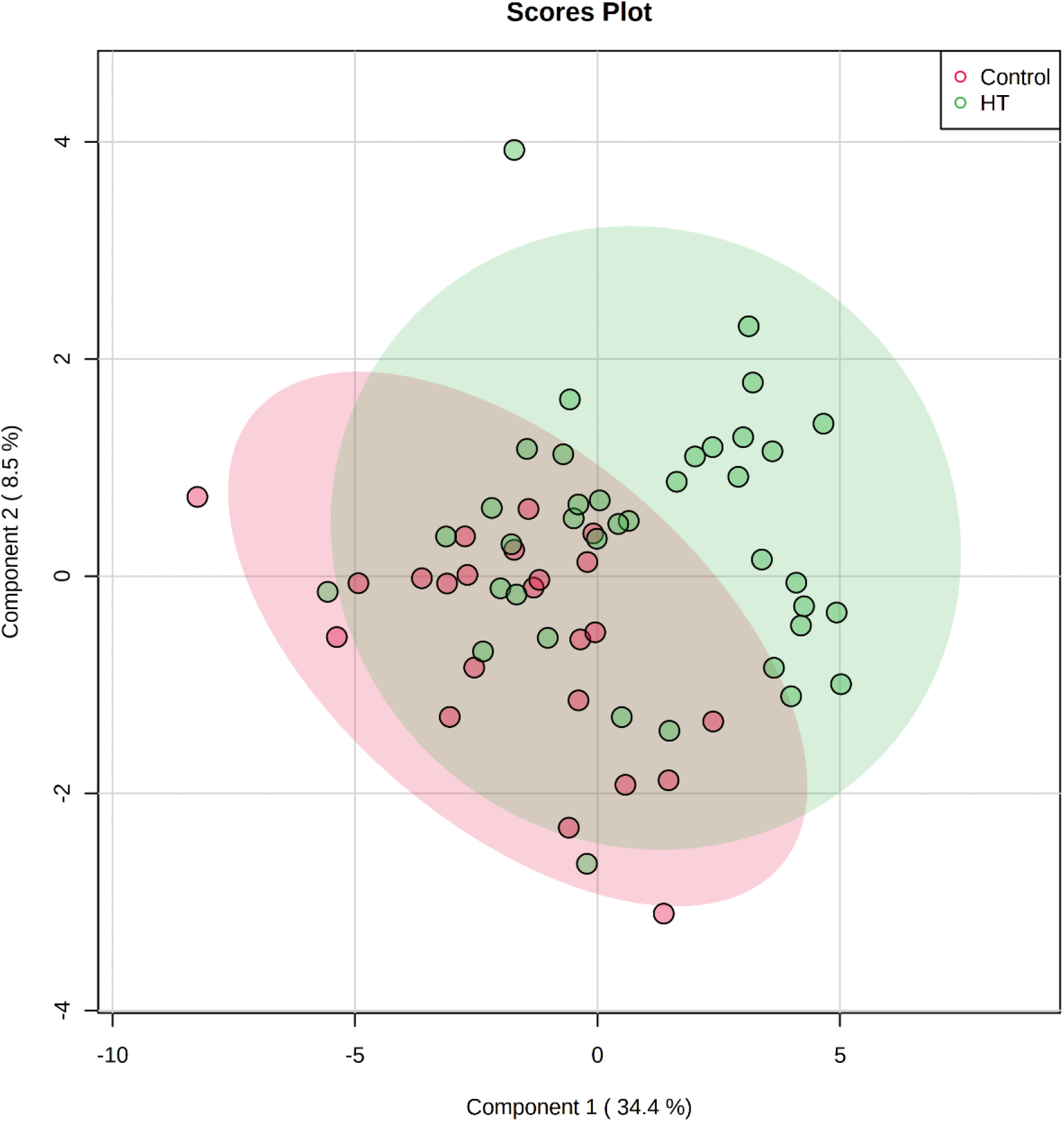
Two-dimensional score plot of follicular fluid samples from the control group (red) and the Hashimoto’s group (HT-green) undergoing in vitro fertilization treatment by PLS-DA. Control n=23. Hashimoto n=38

**Figure 3:**
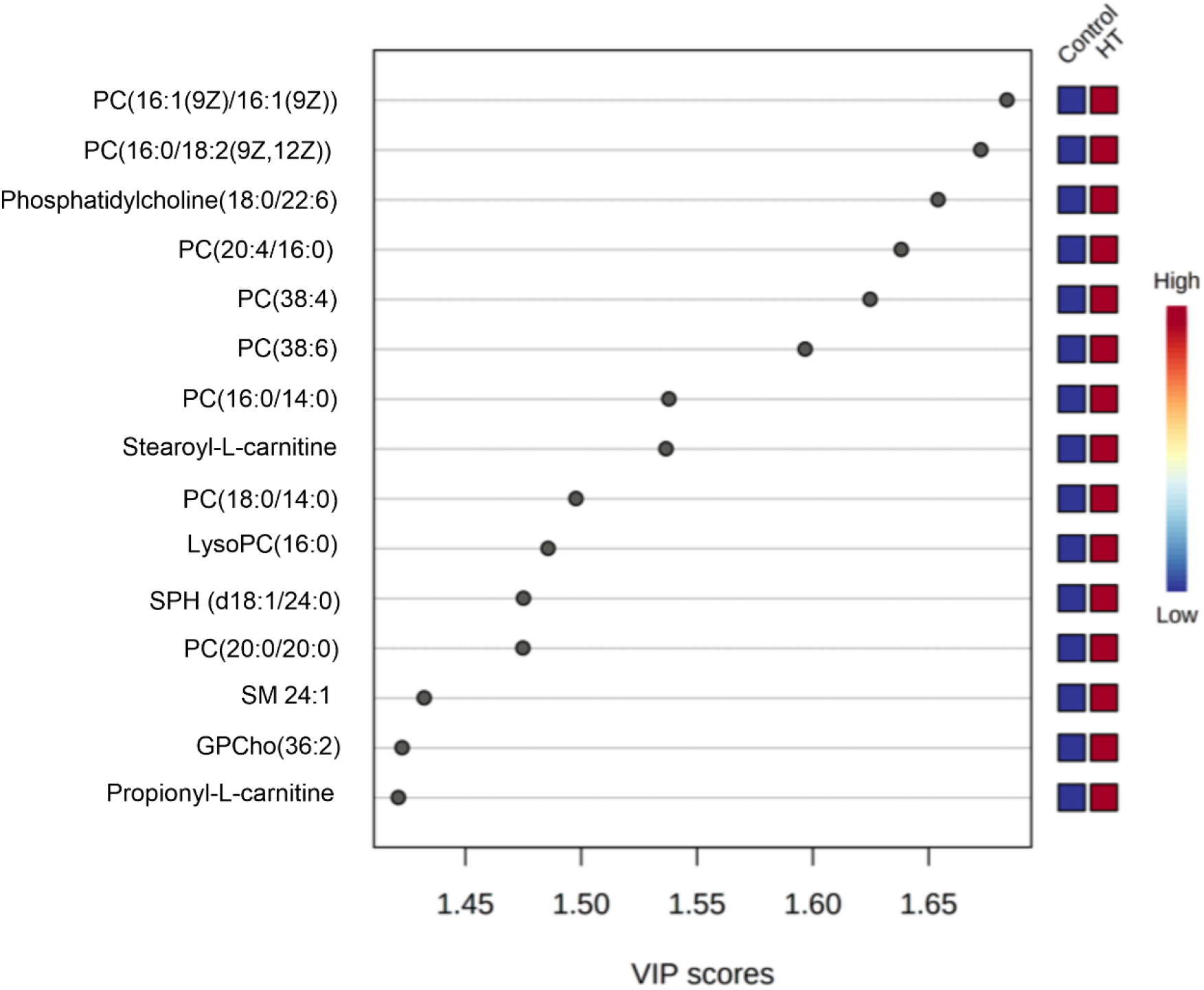
Identification of important metabolite markers among the studied groups. The important features were identified by PLS-DA. The coloured boxes on the right indicate the relative concentrations of the corresponding metabolite in each group. The red-coloured box located below the group indicates that there is a high concentration of the metabolite in this group, while the blue-coloured boxes represent low concentrations. HT: Hashimoto’s group, PC/GPCho: phosphatidylcholine, SPH/SM: sphingomyelin, LysoPC: lysophosphatidylcholine. Control n=23. Hashimoto n=38

**Figure 4:**
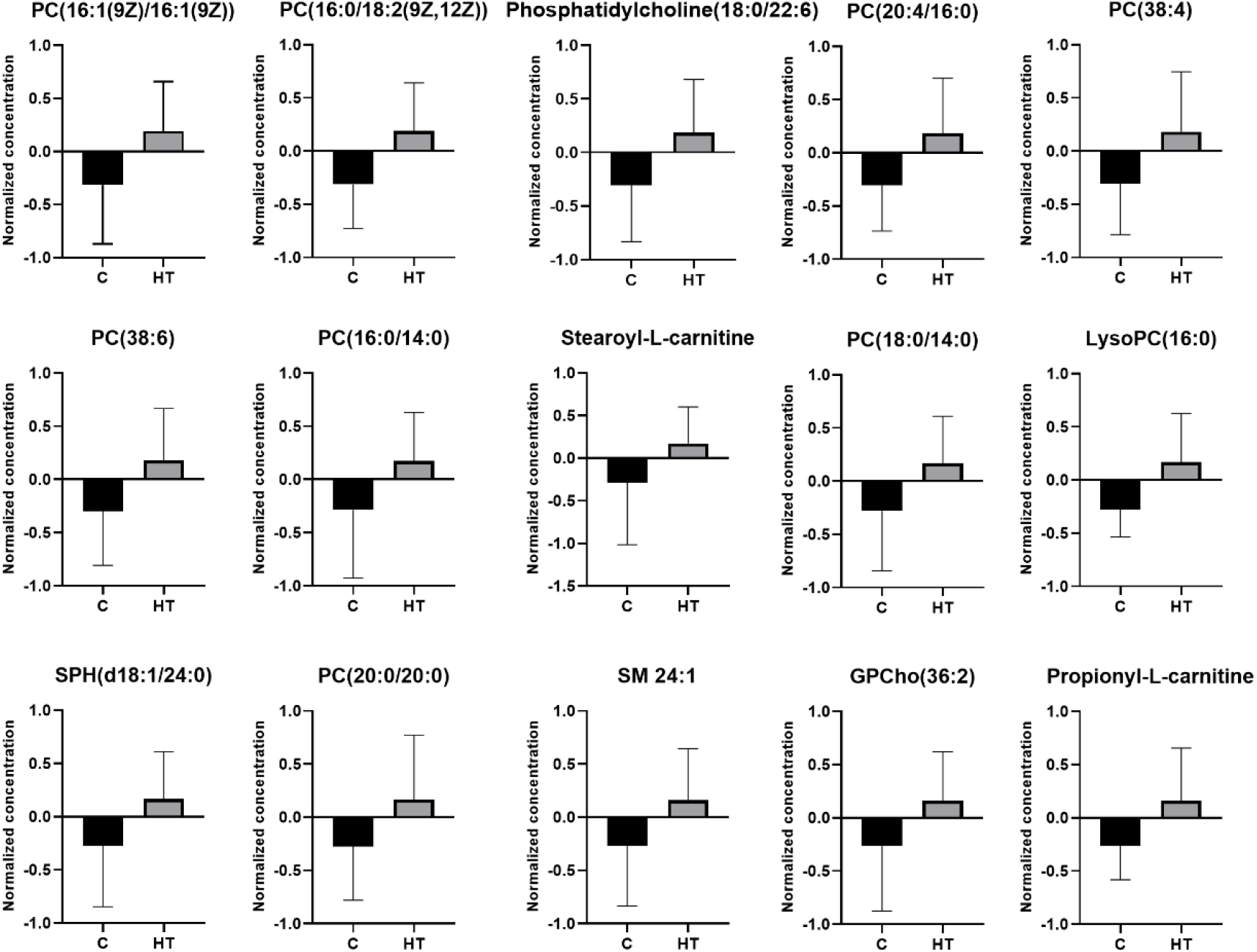
Normalized concentrations of 15 metabolic markers found in the follicular fluid of patients. The red colour represents the control group and the green colour represents the Hashimoto’s group. Data are expressed as mean and standard deviation. HT: Hashimoto’s group, PC/GPCho: phosphatidylcholine, SPH/SM: sphingomyelin, LysoPC: lysophosphatidylcholine. Control n=23. Hashimoto n=38

### 3.4 Pathway analysis

We searched for possible affected pathways based on the identified metabolites and their respective concentrations. We detected five altered pathways: the glycerophospholipid, arachidonic acid, linoleic acid, alpha-linolenic acid, and sphingolipid metabolism pathways (Figure 5, Table 3).

**Figure 5:**
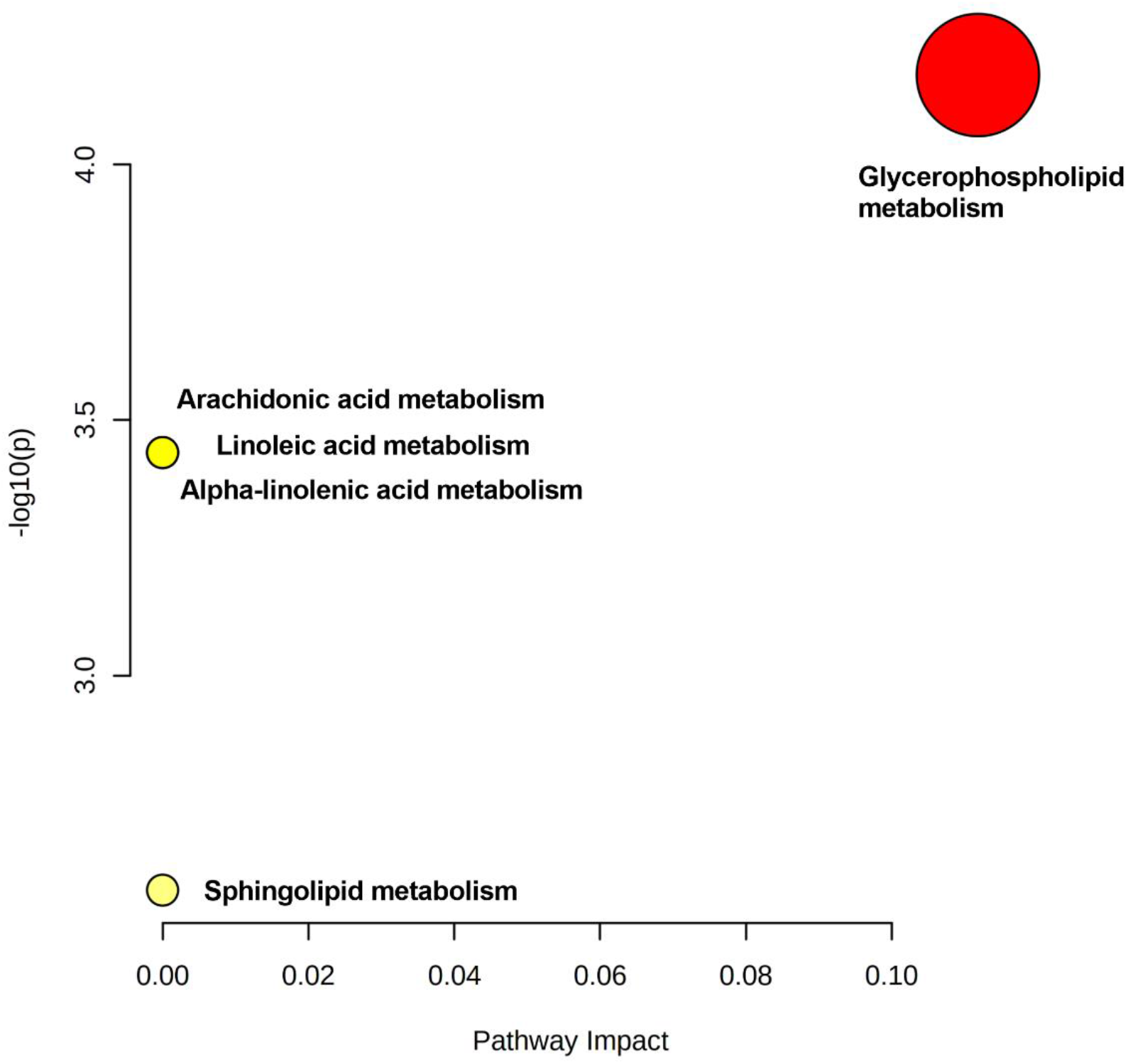
Pathway analysis summary showing altered follicular fluid metabolism of patients with Hashimoto’s thyroiditis. The x-axis means the pathway impact value and the y-axis means the -log of the P-value obtained from pathway enrichment analysis. The size of the circle represents your pathway enrichment and the darker the colors of the pathways the more significance it is.

**Table 3:**
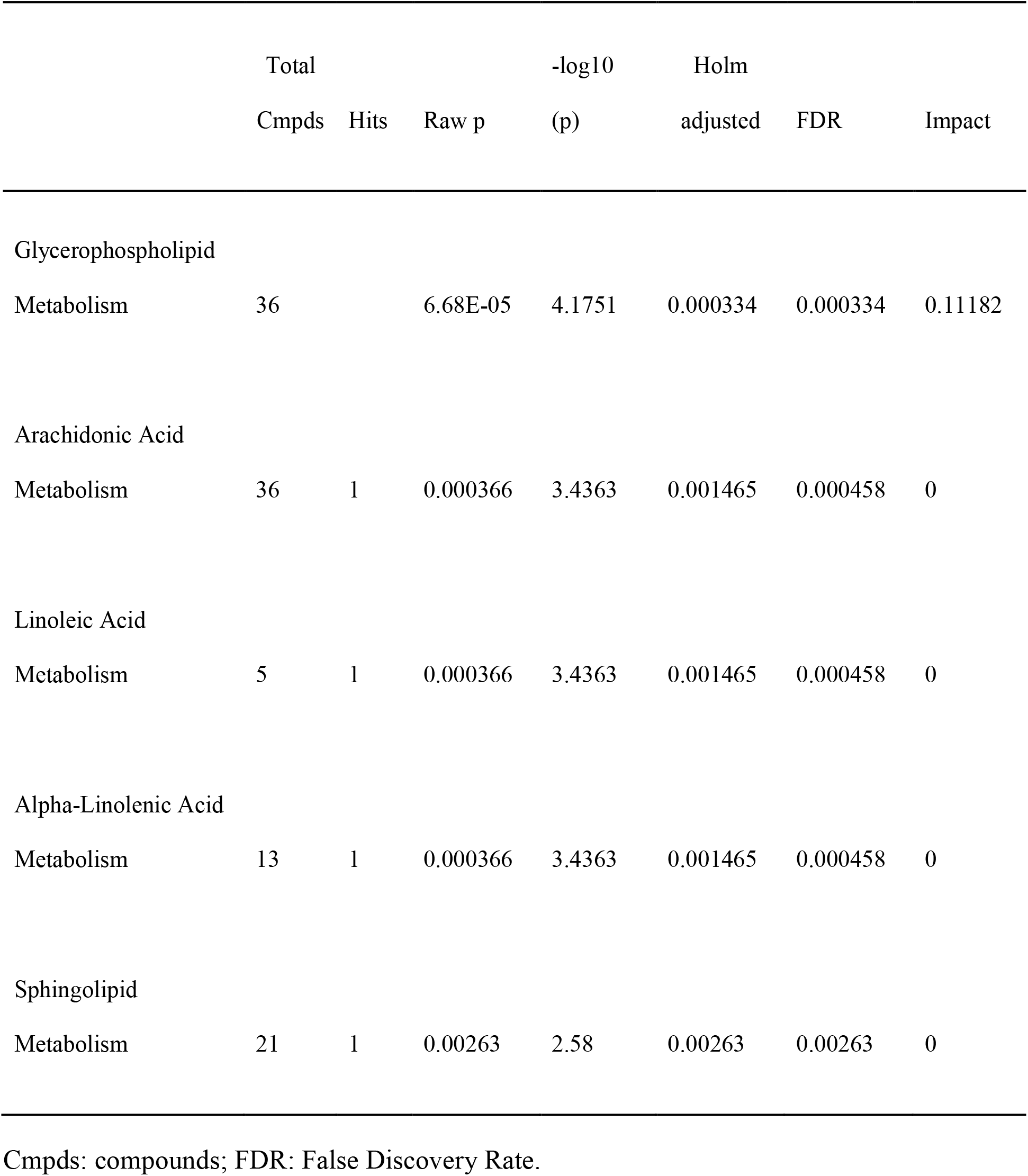
Pathway analysis results based on metabolic markers and their concentrations found in follicular fluid.

## 4 Discussion

The mechanism through which HT affects female fertility is still not known. However, our findings contribute to explaining this connection by showing important differences in the metabolites present in the FF of euthyroid HT patients compared to euthyroid non-HT controls for the first time.

The results of blood measurements of thyroid-related hormones and autoantibodies demonstrate that patients may have normal serum TSH, FT4, and TPOAb levels and still present with the disease, with high TGAb levels. This finding reinforces the importance of continuously measuring autoantibodies, since 5–10% of euthyroid women may have HT (Poppe *et al*., 2021)). Although the measurement of TPOAb is the most used test for HT diagnosis (Alexander *et al*., 2017), our data indicate that serum TGAb levels may be a better marker of the disease. This result conflicts with the current understanding that TPOAb alone would be sufficient to investigate thyroid autoimmunity (Aminorroaya *et al*., 2008)). Although limited to women undergoing IVF, this finding is of clinical relevance since many studies and diagnoses focus only on TPOAb as a diagnostic marker for the disease.

Studies have presented results that stimulate discussion about the clinical relevance of the TGAb dosage (Matalon *et al*., 2003; González *et al*., 2018; Teng *et al*., 2022). A study involving pregnant patients obtained data that contradicted the importance of TGAb dosing. The work showed that there are TPOAb-negative and TGAb-positive patients with normal thyroid function who may need treatment. The article also recommended that TGAb be measured in TPOAb-negative pregnant patients (González *et al*., 2018). Another group showed that pregnant patients with TPOAb positivity and patients with only TGAb positivity had different outcomes for their children (Teng *et al*., 2022). Furthermore, in mice immunized with Tg and presenting high levels of Tg antibodies without thyroid dysfunction, a higher incidence of foetal resorption and reduced placental and embryo weights were observed (Matalon *et al*., 2003).

We found no difference in the evaluated IVF results in relation to the disease. Our findings contrast with previous research showing an increased miscarriage rate and a decreased fertilization rate in women with HT (Monteleone *et al*., 2011; Chen *et al*., 2017; Andrisani *et al*., 2018; Medenica *et al*., 2018; Dong *et al*., 2020); however, our results are in line with other studies suggesting that HT does not affect IVF results (Busnelli *et al*., 2016; Sakar *et al*., 2016; Chen *et al*., 2017; Venables *et al*., 2020). A recent systematic review with meta-analysis suggested that studies evaluating different antibodies (TPOAb and/or TGAb) and different cut-off values may reach conflicting conclusions about the impact of autoimmune thyroiditis on IVF outcomes (Venables *et al*., 2020). Therefore, we believe that our findings alone do not exclude a possible relationship between HT and IVF results.

The FF of HT patients had 15 metabolites with increased concentrations compared to the FF of controls. Our findings included an increase in phosphatidylcholine, acylcarnitine, lysophosphatidylcholine, and sphingolipids. Metabolomics has already been used to investigate HT in blood, urine, and thyroid biopsy samples (Tsoukalas *et al*., 2019, 2020; Capitoli *et al*., 2020; J. Liu *et al*., 2020; Piras *et al*., 2021), but the metabolites found to be altered in these samples were different from those identified in ovarian FF in the present study. However, one of the metabolites that we found to be high in the FF of HT patients, PC (18:0/22:6), was reported to be increased in the blood of hyperthyroid patients (J. Liu *et al*., 2020). Therefore, this is the second study that relates PC (18:0/22:6) with thyroid pathology, and further studies are needed to investigate the possible relationship.

The larger metabolite group in the FF of HT patients was phosphatidylcholine. We identified ten phosphatidylcholines: PC(16:1(9Z)/16:1(9Z)), PC(16:0/18:2(9Z,12Z)), phosphatidylcholine(18:0/22:6), PC(20:4/16:0), PC(38:4), PC(38:6), PC(16:0/14:0), PC (18:0/14:0), PC(20:0/20:0), and GPCho(36:2). Phosphatidylcholine is the lipid subclass that is most abundant in cell membranes. In porcine FF, phosphatidylcholine was found in higher concentrations in large antral follicles than in smaller antral follicles (Lai *et al*., 2018). These data indicate that the synthesis of these metabolites may be involved in follicle growth and oocyte maturation. Furthermore, increased levels of phosphatidylcholine were found in serum and FF samples from patients with ovarian endometriosis (Vouk *et al*., 2012; Cordeiro *et al*., 2015). Therefore, phosphatidylcholine appears to play an important role in ovarian physiology.

One of the metabolites we found to be differentially concentrated in FF from HT patients was a lysophosphatidylcholine, LycoPC(16:0). Lysophosphatidylcholine is secreted by cumulus cells (Gómez-Torres *et al*., 2015) and can induce an acrosome reaction in human sperm (Byrd and Wolf, 1986). The acrosome reaction is necessary for sperm to be able to fertilize an oocyte. The use of a high concentration of this metabolite was able to induce an acrosome reaction in human sperm within 15 minutes. However, this same experiment also identified that sperm had a rapid loss of motility and a drop in viability (Byrd and Wolf, 1986). The increased concentration of lysophosphatidylcholine in FF, which is released along with the oocyte at the time of ovulation, can decrease sperm motility and viability during fertilization in vivo. If the increase in FF lipid concentration can have adverse effects on sperm (Byrd and Wolf, 1986), this could explain the difficulty that patients with HT experience in becoming pregnant naturally and why our study did not identify such a gap in pregnancy rates following IVF treatment. In IVF, the oocytes are removed from the FF and kept in culture medium, so the composition of the FF does not affect the sperm. Further experimental studies in suitable animal models are needed to directly test the hypothesis that increasing the concentration of lysophosphatidylcholine in FF can hinder in vivo fertilization.

We found an increase in acylcarnitines, propionyl-L-carnitine, and stearoyl-L-carnitine in the FF of women with HT. Acylcarnitine is responsible for carrying out the beta-oxidation of fatty acids. This process is one of the most important pathways for producing metabolic energy (Indiveri *et al*., 2011). Acylcarnitine supplementation is protective against mouse oocyte cytoskeletal damage and embryonic apoptosis induced by incubation in the peritoneal fluid of patients with endometriosis (Mansour *et al*., 2009). Furthermore, this supplementation in mice increased beta-oxidation and improved the rate of fertilization and the development of embryos (Dunning *et al*., 2011). In addition to findings in animal models, acylcarnitine has also been shown to be important in studies involving humans. A reduction in acylcarnitine levels has been previously noted in serum and FF samples from patients who had more than 9 oocytes and more than 6 embryos (Várnagy *et al*., 2013). The study suggests that in patients with better reproductive potential, this pathway is upregulated and, therefore, there is high consumption of acylcarnitine, with its concentrations found at low levels (Várnagy *et al*., 2013). Based on this work, it is possible to relate the increase in acylcarnitine levels in the FF of women with HT compared to control women with low consumption of this metabolite in our study, providing a further potential mechanism for the subfertility associated with HT, which can be overcome by IVF. More work is needed to investigate the relationship of acylcarnitine with reproductive potential and disease.

Sphingomyelins are a type of sphingolipid found in animal cell membranes. Here, we found an association between HT and increased FF levels of sphingomyelins SM 24:1 and SPH (d18:1/24:0), which are important components of the sphingomyelin pathway. In addition, sphingolipids have been linked to ovarian cancer and ovarian endometriosis (Vouk *et al*., 2012; Cordeiro *et al*., 2015; Zeleznik *et al*., 2019). An increase in this metabolite in the blood has been associated with a risk of developing ovarian cancer 3 to 23 years before diagnosis (Zeleznik *et al*., 2019).

The metabolic changes found in FF suggest changes in five pathways, namely, the glycerophospholipid, arachidonic acid, linoleic acid, alpha-linoleic acid, and sphingolipid pathways. The first involves phosphatidylcholine and lysophosphatidylcholine. This was previously reported to be dysregulated in patients with ovarian endometriosis (Xu *et al*., 2018). Arachidonic acid and linoleic acid derivatives play relevant roles that can affect fertility and pregnancy. Some of its derivatives may be predictive markers of pregnancy complications such as gestational diabetes mellitus or preeclampsia (Szczuko *et al*., 2020).

In the sphingolipid pathway, there are metabolites related to the apoptosis process, such as ceramide and sphingosine, and metabolites related to cell survival in response to apoptotic stimuli, such as sphingosine-1P (Cuvillier, 2002). Thyroid hormones, mainly T3, induce the survival of human granulosa cells, both in the preovulatory stage and in the luteinized phase (Falzacappa *et al*., 2009; di Paolo *et al*., 2020). A picture of hypothyroidism, which occurs in HT when untreated, could lead granulosa cells to apoptosis due to the increase in the sphingolipid pathway and the decrease in T3.

All our findings involving the metabolomics of FF seem to be directly involved in ovarian physiology and female fertility, although we did not find any significant impact on the patients’ IVF results. This suggests that HT may affect follicle development and oocyte fertilization to an extent that can be bypassed by IVF through ovarian stimulation and in vitro procedures. However, this study has some limitations. We first used metabolomics as an exploratory strategy to identify potential pathways in ovarian FF affected by HT, and our results should be interpreted as hypothesis-generating rather than the conclusive demonstration of such mechanisms. The relationship of FF metabolites and their pathways with ovarian physiology and natural fertility should be further studied in animal models due to ethical constraints in obtaining this biological fluid in women who are trying to conceive spontaneously. In the context of IVF, our findings may direct hypothesis-driven studies to clarify whether the presence of HT has any implication for gamete and embryo development beyond the fertilization rate and gross embryo morphology.

## 5 Conclusions

In this study, we observed that the serum TGAb level may be a more consistent biomarker of HT in women preparing for IVF than the serum TPOAb level, which is the standard HT marker used in clinical practice. For the first time, metabolomics was used to assess the metabolic profile present in the FF of women with the autoimmune disease, and data showed that HT can change the metabolic profile of FF. Furthermore, these findings may explain, how the presence of autoantibodies in FF can impact ovarian follicles and female fertility.

## 8 Acknowledgements

We thank everyone involved who made this work possible.

